# Stress-induced myonectin improves glucose homeostasis by inhibiting glycemic response to HPA axis

**DOI:** 10.1101/838003

**Authors:** Zhengtang Qi, Jie Xia, Xiangli Xue, Jiatong Liu, Xue Zhang, Xingtian Li, Wenbin Liu, Lu Cao, Lingxia Li, Zhiming Cui, Zhuochun Huang, Benlong Ji, Qiang Zhang, Shuzhe Ding, Weina Liu

## Abstract

Inhibiting glycemic response to HPA axis contributes to glycemic control for diabetic patients. Here, mice were subjected to high-fat diet and intermittent chronic stress, and glucose homeostasis and lipolysis were determined during the intervention. Firstly, we found that glucose intolerance appears at the earliest, followed by reduced insulin sensitivity and increased epinephrine (EPI) sensitivity in the early stage of diet-induced obesity. Next we investigated whether chronic stress impairs glycemic control and which mediates its effects. Short-term stress training raises serum and skeletal muscle myonectin (Myn) levels and improves glucose intolerance. Stress attenuates blood glucose and glycerol responses to EPI, but enhances lipolytic response to EPI in adipose tissues. Myn overexpression in vivo improves glucose tolerance and enhances insulin sensitivity at the cost of blunting glycemic responses to EPI. Myn knockdown reduces beneficial effects of stress or exercise on glucose homeostasis. Together, myonectin is a stress-induced myokine that readjusts glycemic and metabolic responses to HPA axis, and thus prevent the progression of glucose intolerance and obesity.

**Graphical Abstract:** 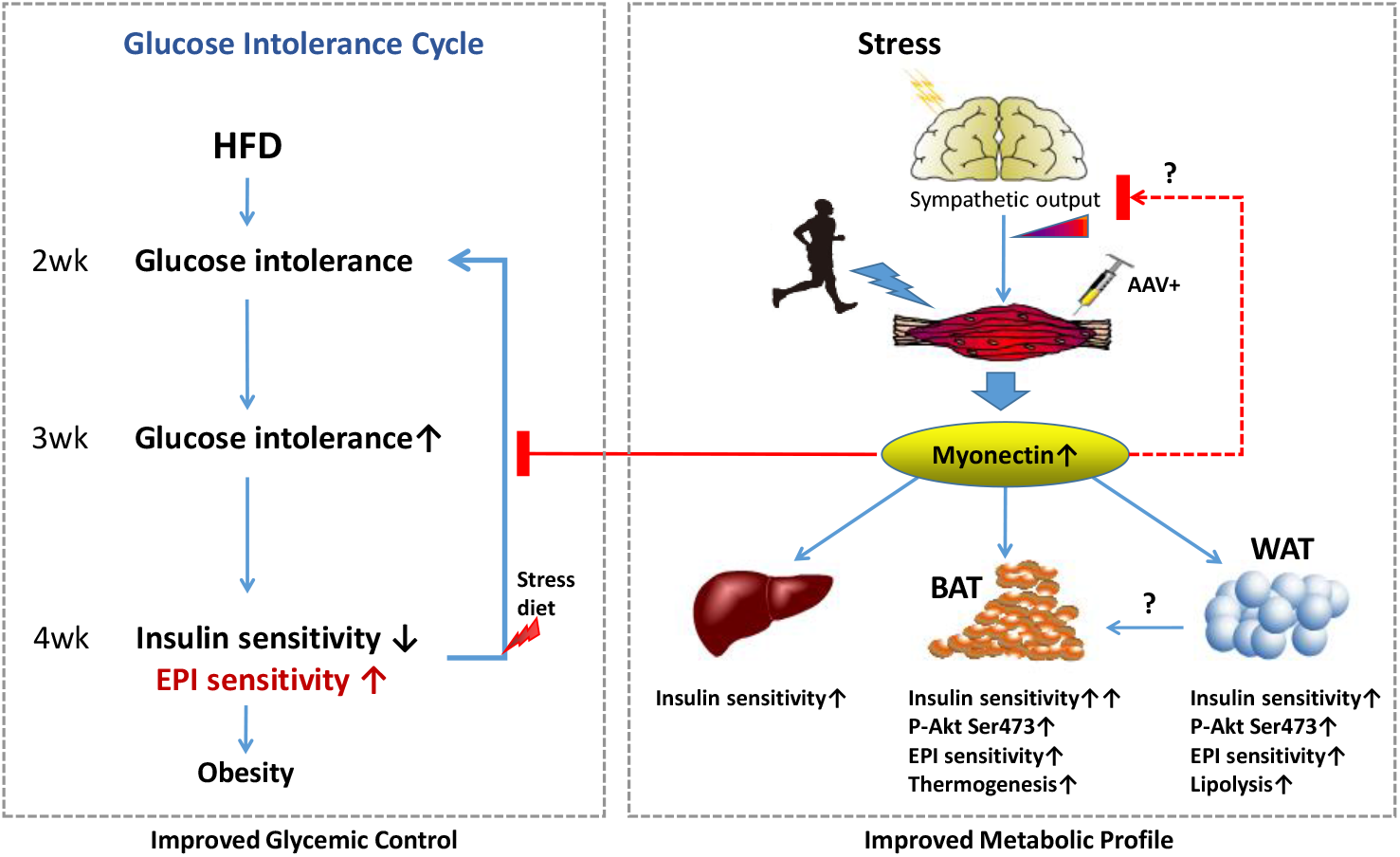

**One Sentence Summary:** Chronic stress breaks glucose intolerance cycle to resist diet-induced obesity, through myonectin-mediated inhibition of glycemic response to epinephrine (EPI) and activation of insulin signaling in adipose tissues.

**Highlights:**

- EPI sensitivity increases after glucose intolerance and with reduced insulin sensitivity in diet-induced obesity
- Chronic stress blunts glycemic responses to EPI and increases myonectin levels in serum and skeletal muscle
- Myonectin attenuates glycemic response to EPI and improves metabolic profile in HFD-fed mice
- Reducing myonectin reverses beneficial effects of stress on glucose homeostasis

## Introduction

Hyperglycemia and diabetes are associated with the activation of hypothalamic-pituitary-adrenal (HPA) axis(Bruehl et al, 2007). Recent studies have demonstrated that suppression of the HPA axis improves glycemic control and reverses diabetes(Perry et al, 2015; Perry et al, 2014). Chronic stress and exercise are both important exogenous factors to induce the activation of HPA axis. Chronic stress increased sympathetic nervous activity and elevated basal corticosterone levels and rapid glucocorticoid production, so stress exposure increases blood glucose via the activation of HPA axis (Lowrance et al, 2016). However, chronic exercise enhanced HPA-axis negative feedback and thus reduced HPA responses to stress or next bout of exercise (Hare et al, 2014). Thus, chronic stress is associated with a higher risk of type 2 diabetes(Heraclides et al, 2009), while regular exercise is helpful for glycemic control in the previous cohort studies(Lloyd & Barnett, 2008). High-fat feeding enhances HPA axis tone in a manner that resembles sympathetic overdrive in chronic stress exposure (McNeilly et al, 2015; Ryan et al, 2018). Thus, lowering glycemic response to HPA axis by some means contributes to glycemic control. In addition, recent studies focusing on the impact of stress on obesity suggest two conflicting conceptions: 1) stress promotes obesity; 2) stress results in weight loss or obesity resistance. Diet and brown fat lead to the dichotomous effect of stress on metabolic outcomes (Razzoli & Bartolomucci, 2016). This contradiction implies that adipose tissues can respond differently to HPA axis hyperactivity in regulating lipid metabolism.

Stable blood glucose control is usually the result of the integration of the central nervous system with multiple peripheral organs. Studies showed that adipose tissue, skeletal muscle and liver are psychosocial stress-responsive tissues, making an important understanding of the central nervous circuits linking stress coping with energy homeostasis. Psychological stress drives the sympathetic thermogenesis in brown adipose tissue (BAT) and contributes to hyperthermia (Kataoka et al, 2014). Subordination stress selectively inhibits the insulin signaling pathway in liver and skeletal muscle. Conversely, white adipose tissue (WAT) shows increased insulin sensitivity and enhanced fat deposition (Sanghez et al, 2016). Chronic psychosocial defeat stress differently affects lipid metabolism in liver and WAT and induces hepatic oxidative stress in HFD mice (Giudetti et al, 2019). These findings suggest that different tissues respond differently to stress in insulin signaling, lipid metabolism, and thermogenesis, eventually leading to a metabolic profile of the whole-body. The metabolic phenotype under stress should be an outcome of reciprocal regulation between brain and peripheral tissues. For example, the dorsomedial hypothalamus functions as a hub for stress signaling and mediates a psychosocial stress-induced thermogenesis in BAT (Kataoka et al, 2014). There is still a need for exploring the mediator that reciprocally regulate glucose and lipid metabolism between brain and peripheral tissues.

Chronic unpredictable mild stress (CUMS) is used to induce depression-like behaviors in rodents (Antoniuk et al, 2018). In addition, CUMS often leads to weight loss, suggesting that CUMS has the potential to prevent obesity. Studies have shown that chronic stress aggravated glucose intolerance but not obesity in db/db mice on the basis of a disrupted leptin circuitry (Razzoli et al, 2015). Chronic stress enhanced human vulnerability to diet-related metabolic risk (Aschbacher et al, 2014). There is also positive evidence that chronic variable stress improved glucose tolerance in rats with prediabetes (Packard et al, 2014). Chronic social defeat stress induced hypophagia and weight loss, ultimately improving glucose tolerance and serum levels of insulin and leptin in diet-induced obesity (Balsevich et al, 2014). Together, it is difficult to draw a clear conclusion on the effect of chronic stress on metabolic outcomes, due to the differences in stress mode (variable or invariable), pathogenic model (dietary or genetic), duration of sustained stress (short-term or long-term), and research object (animal or human) in the previous studies. In this study, we are concerned about the early process of diet-induced obesity involving elevated sympathetic activity, so we do not expect stress to cause depressive behaviors in mice. To further explore the evidence of stress controlling dietary obesity, we treated mice with short-term dietary intervention and stress training.

Recently, myonectin (Myn) has been identified as a myokine that links skeletal muscle to systemic lipid homeostasis and liver autophagy, and predicts the development of type 2 diabetes (Li et al, 2018; Seldin et al, 2013; Seldin et al, 2012). Moreover, Myn functions as an exercise-induced myokine which ameliorates myocardial ischemic injury by suppressing apoptosis and inflammation in heart (Otaka et al, 2018). Myn expression is stimulated by EPI and the activation of cAMP-PKA signaling (Seldin et al, 2012), so we will explore the role of Myn in stress-induced weight loss. Our findings reveal that stress training promotes the expression and secretion of Myn, mediating to improve glucose intolerance, increase BAT thermogenesis, and enhance insulin and EPI sensitivity in adipose tissue. This improvement comes at the cost of EPI resistance and blunting glycemic response to stress. We identified Myn as a stress-responsive myokine that mediates sympathetic activity to maintain glycemic control and protects against diet-induced obesity.

## Results

### Enhanced EPI sensitivity appears after glucose intolerance and induces glucose intolerance cycle in the early stage of diet-induced obesity

To understand the longitudinal change in glucose homeostasis in the development of diet-induced obesity, we performed glucose, insulin and EPI tolerance test in HFD-fed mice every week after diet intervention. At the 2^nd^ week, glucose intolerance emerged firstly in HFD-fed mice, but the difference was not found in insulin and EPI tolerance test (Fig. 1A). The results in week 3 were the same as those in week 2 (Fig. S1A). At the 4^th^ week, glucose intolerance was further aggravated; moreover, IPITT showed that HFD have led to insulin resistance in mice (Fig. 1B). Blood glucose increased within 30 min after EPI injection and then began to fall back in normal chow-fed mice, but blood glucose increased for 60 min before it began to fall back in the HFD-fed mice (Fig. 1B, D). This suggests that HFD increased the glycemic response of the mice to EPI. Our data suggest that HFD induces an impairment of glucose homeostasis, followed by insulin resistance and EPI hypersensitivity. Elevated EPI sensitivity may further aggravate glucose intolerance in mice exposed to sympathetic activity, such as chronic stress.

**Fig.1.**
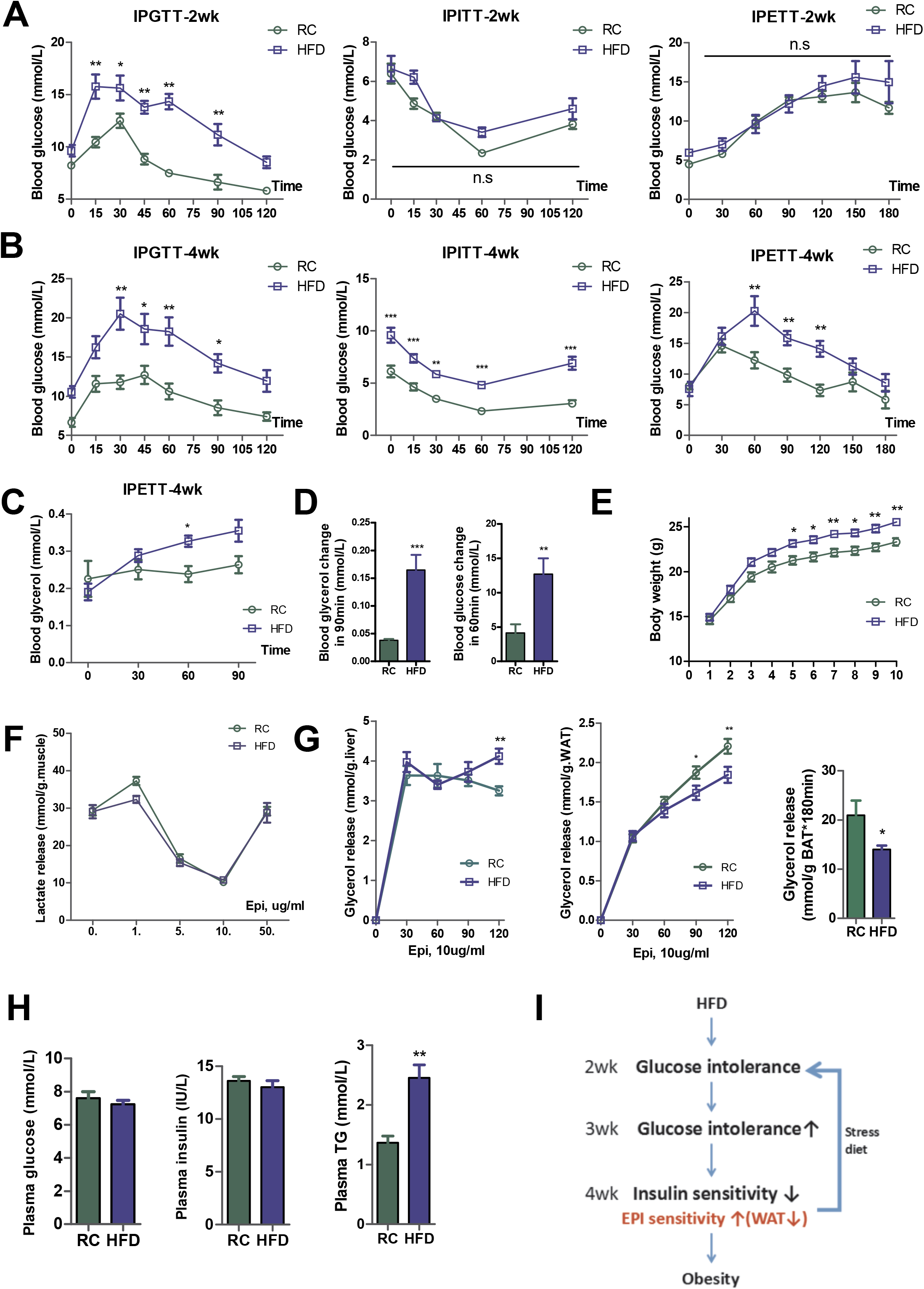
Increased EPI sensitivity occurs after glucose intolerance and concurrently with insulin resistance in the early stage of diet-induced obesity. (A) Glucose tolerance tests (IPGTT), Insulin tolerance tests (IPITT) and Epinephrine tolerance tests (IPETT) at the 2nd week of dietary intervention. n=6–7/group. (B) Glucose tolerance tests (IPGTT), Insulin tolerance tests (IPITT) and Epinephrine tolerance tests (IPETT) at the 4th week of dietary intervention. n=6–7/group. (C) Blood glycerol levels in the IPETT at the 4^th^ week. n=6–7/group. (D) Blood glycerol changes in 90 min and blood glucose changes in 60 min after EPI injection in the 4^th^ week IPETT. n=6–7/group. (E) Body weight of mice treated with regular chow and HFD. n=8/group. (F) EPI-stimulated lactate production in the cultured gastrocnemius muscles. n = 6/group at each concentration point of EPI. (G) EPI-stimulated glycerol release in 120~180 min after EPI was added into the cultured liver tissues, epididymal white adipose tissue (eWAT) and brown adipose tissue (BAT). n=6–7/group. (H) Plasma glucose, triglyceride (TG) and insulin levels. n = 14–16/group. (I) Schematic summary of glucose intolerance cycle that promotes diet-induced obesity. RC, regular chow; HFD, high-fat diet. Statistical significance was evaluated by unpaired two-tailed Student’s t test or two-way ANOVA with Tukey post hoc tests. Data are means ± SEM. *p < 0.05, **p < 0.01, ***p < 0.01 versus RC.

Childhood obesity is associated with sympathetic overactivity and EPI resistance in the lipolysis of adipose tissue (Bougneres et al, 1997; Sorof & Daniels, 2002). Since EPI can lead to fat mobilization, we also evaluated the changes in serum glycerol after intraperitoneal injection of EPI. Our results showed that little change was found in serum glycerol within 90min after EPI injection in normal diet mice, but serum glycerol increased significantly in HFD mice (Fig. 1C, D). This suggests that HFD increases the sensitivity of fat mobilization to EPI. HFD resulted in significant weight gain from week 3 (weighing twice a week, Fig. 1E). HE staining of adipose tissue showed a marked increase in adipocyte size (Fig. 1F). These data suggest that 4-week HFD led to an early phenotype of obesity in mice with increased EPI sensitivity. This phenotype is partly consistent with the particularity of childhood obesity.

We also tested the EPI sensitivity of skeletal muscle, liver, and white adipose tissue in vitro after the sacrifice of mice. In dose-dependent experiments on skeletal muscle culture, the release of lactic acid from cultured gastrocnemius increased at 1ug/ml EPI, decreased at 1-10ug/ml EPI, and increased sharply at 50ug/ml EPI (Fig. 1G). This curve suggests that skeletal muscles respond differently to different concentrations of EPI. There are types of adrenergic receptors (ARs) on the cell surface, which respectively mediate lipolytic or antilipolytic pathways in human adipose tissue (Stich et al, 2002). In obese men, exercise increases sympathetic and adrenergic activity, but induces a desensitization in β1- and β2- adrenergic lipolytic pathways and the stimulation of α2ARs- mediated antilipolytic action in subcutaneous adipose tissue (Marion-Latard et al, 2001; Stich et al, 2000). This suggests that obese people can initiate fat protection during exercise by modulating the activity of ARs. Our results suggest that types of ARs may also exist in skeletal muscle in response to different levels of adrenergic stimulation, but we did not find a difference in skeletal muscle EPI sensitivity with HFD.

In the time-dependent experiment, we measured the release of glycerol into the culture medium during the incubation of liver, WAT and BAT with 10ug/ml EPI. Our results showed that the liver and WAT glycerol release increased rapidly within the first 30 minutes, but the liver of HFD mice released more glycerol and WAT released less glycerol at the end of 120 min incubation (Fig. 1H). This suggests that HFD leads to opposite changes in the EPI sensitivity of liver and WAT. This change promoted the liver to remove fat and reduced the lipolysis of WAT, suggesting the capability of sympathetic activity to transfer fat from liver to WAT. Previous studies showed that diet-induced obesity increased inflammatory macrophages counts in adipose tissue rather than the liver. This was associated with greater fat gain in adipose tissue compared with liver (de Meijer et al, 2012). Here, we did not found that 4-week HFD resulted in fat deposition in the liver (Fig. S1B). HFD increased expression of the adrenergic receptors in WAT, but reduced expression of Adrb1, Adrb3, HSL and Rgs2 in BAT (Fig. S1C). Our results suggest that tissue specificity in EPI sensitivity is responsible for greater fat gain in adipose tissue. In tissues culture with 180 min EPI incubation at 10ug/ml, glycerol release showed that HFD impaired the EPI sensitivity of BAT (Fig. 1I).

In addition, HFD led to an increase in serum triglycerides, but no significant changes in serum levels of glucose, insulin, leptin, glycerol, EPI and myonectin (Fig. 1J, S1F). Together, HFD leads first to glucose intolerance and secondly to insulin resistance and increased EPI sensitivity in the development of diet-induced obesity. However, adipose tissue lipolysis is less sensitive to EPI. We believe that increased sensitivity of blood glucose to EPI and decreased sensitivity of fat lipolysis to EPI will further promote glucose intolerance and fat deposition in the diet-induced obesity (Fig. 1K).

### Stress training blunts glycemic response to EPI and enhances Myn levels in circulation and skeletal muscle

Neuroendocrine studies demonstrated HPA axis overactivity in CUMS-induced depression(Keller et al, 2017), suggesting a role of excessive EPI release in enhancing energy expenditure and thus preventing obesity. In this study, 3 weeks of stress training prevented diet-induced obesity (Fig. 2D, E), without depression and anxiety-like behaviors in mice (Fig. S2A-D). Stress training improved glucose intolerance, but insulin sensitivity was not altered (Fig. 2A). Glycemic response to EPI was attenuated in the stressed mice (Fig. 2B). Although stress training increased the basal level of serum glycerol, there was no significant fluctuation of serum glycerol level within 90min after EPI injection (Fig. 2C). Stress training increased serum insulin and glycerol levels, but decreased serum triglyceride levels (Fig. 2H). These results indicate that stress training enhanced the ability of mice to maintain metabolic homeostasis, and improved metabolic profile in HFD-fed mice.

**Fig.2.**
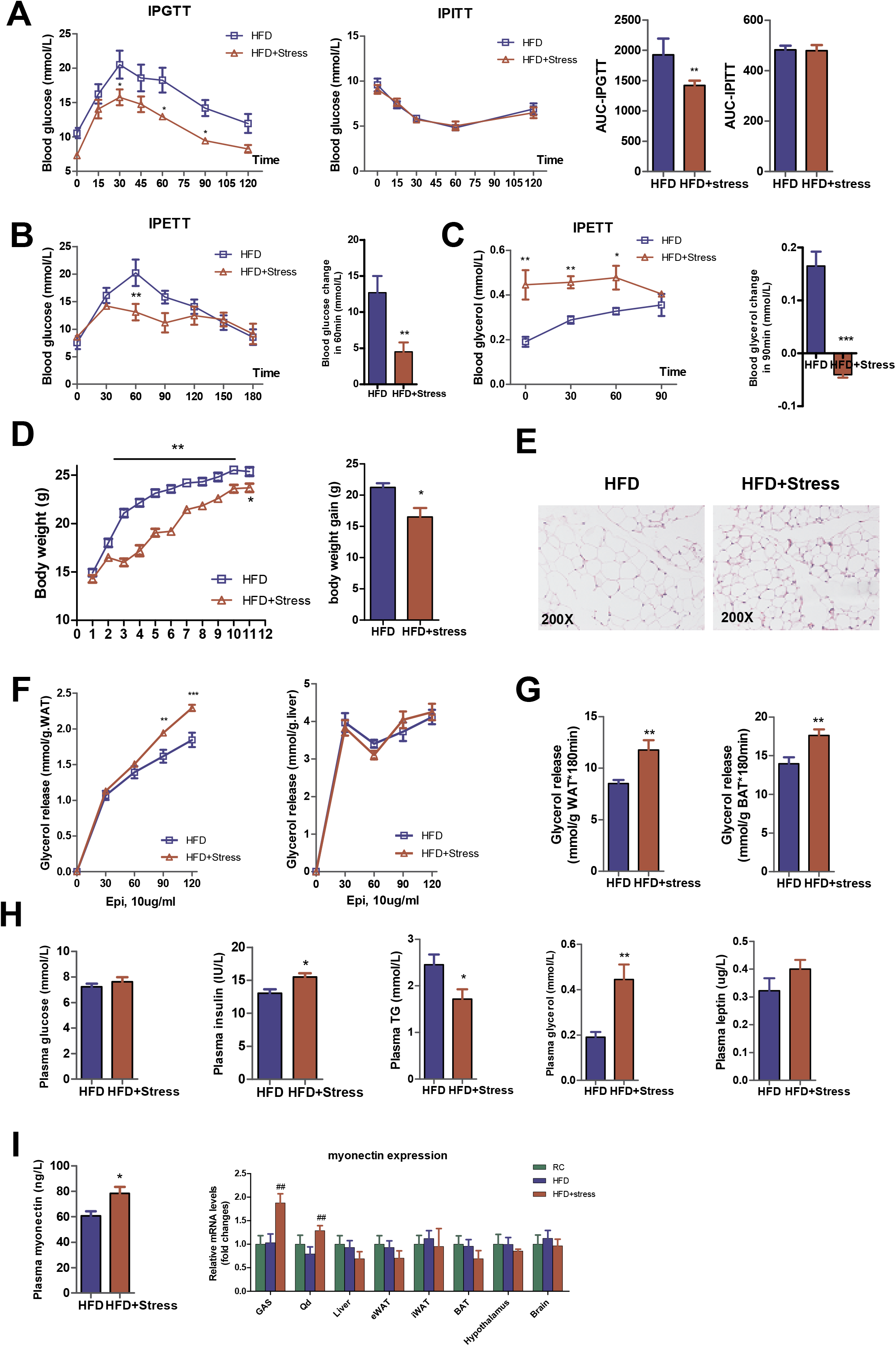
Stress training improves glucose intolerance and reprograms glycemic and lipolytic responses to EPI in HFD-fed mice. (A) Glucose and insulin tolerance tests in HFD-fed mice treated with or without stress training. Quantitative values are the area under the curve (AUC) ± SEM. n = 6–7/group. (B) Epinephrine tolerance tests (IPETT) and blood glucose changes in 60 min after EPI injection. n = 6–7/group. (C) Epinephrine tolerance tests (IPETT) and blood glycerol changes in 90 min after EPI injection. n = 6–7/group. (D) Body weight and body weight gain of mice. n = 8/group. (E) Representative H&E stained eWAT. Original magnification is × 200. n = 6/group. (F) EPI-stimulated glycerol release in 120 min after EPI was added into the cultured tissues of liver and eWAT. n = 6/group at each time point. (G) EPI-stimulated glycerol release in 180min after EPI was added into the cultured tissues of eWAT and BAT. n = 6/group. (H) Plasma glucose, insulin, triglyceride (TG), glycerol and leptin levels. n = 10–12/group. (I) Plasma myonectin levels and myonectin expression in various tissues. ##p < 0.01 versus HFD. Statistical significance was evaluated by unpaired two-tailed Student’s t test or two-way ANOVA with Tukey post hoc tests. Data are means ± SEM. *p < 0.05, **p < 0.01, ***p < 0.001 versus HFD.

Tissues incubated with EPI in vitro confirmed that stress training increased the sensitivity of WAT and BAT to EPI, compared with the HFD control group (Fig. 2F, G). Although we did not observe a significant improvement in systemic insulin sensitivity, insulin-stimulated glucose uptake was increased in skeletal muscle of stressed mice (Fig. S2G). EPI stimulates the expression of Myn, while HFD reduce Myn expression in myotubes (Seldin et al, 2012). Stressed mice fed HFD had less adipose tissue as well as lower serum leptin levels. Inhibition of beta(3)-adrenergic signaling during social stress improved these metabolic abnormalities but worsens behavioral symptoms (Chuang et al, 2010). Although the results of this study are not completely consistent with our data, it suggests that social or psychological stress may regulate energy metabolism and adipose tissue homeostasis through beta(3)-adrenergic signaling. As an EPI-induced myokine, myonectin is likely to be released from muscles as a byproduct of stress - sympathetic excitation. As we expect, stress training increased serum Myn levels and Myn expression in gastrocnemius and quadriceps muscles (Fig. 2I). This suggests that Myn may be a stress-responsive myokine that mediates stress to improve metabolic profiling. Together, our results indicate that stress training may not cause depressive-like behavior, but increases the expression and secretion of Myn in skeletal muscle and thus improves metabolic profile in HFD-fed mice.

### Induction of sympathetic activity enhances myonectin expression in skeletal muscle

To further establish the link between sympathetic activity and myonectin in skeletal muscle, we asked whether sympathetic activity could enhance myonectin expression. In an independent study, we found that Myn expression was up-regulated in skeletal muscle immediately after running exercise, but Myn expression was inhibited due to excessive exercise intensity (Fig. 3A). Serum Myn levels increased after acute exercise but not chronic exercise training (Fig. 3B). We further detected Myn expression in cultured muscles stimulated by EPI at different concentrations. Lower concentrations of EPI induced an increase in Myn expression, but higher concentrations of EPI inhibited Myn expression. The expression of Adrb3 increased at lower EPI, but decreased at higher EPI (Fig. 3C). This suggests that skeletal muscles switch off some adrenergic signaling at higher sympathetic stimuli to avoid stress injury. Myn expression in myotubes is up-regulated by increase in cellular cAMP or calcium levels (Seldin et al, 2012), so we hypothesized that Adrb3 mediates EPI-induced Myn expression in skeletal muscle.

**Fig.3.**
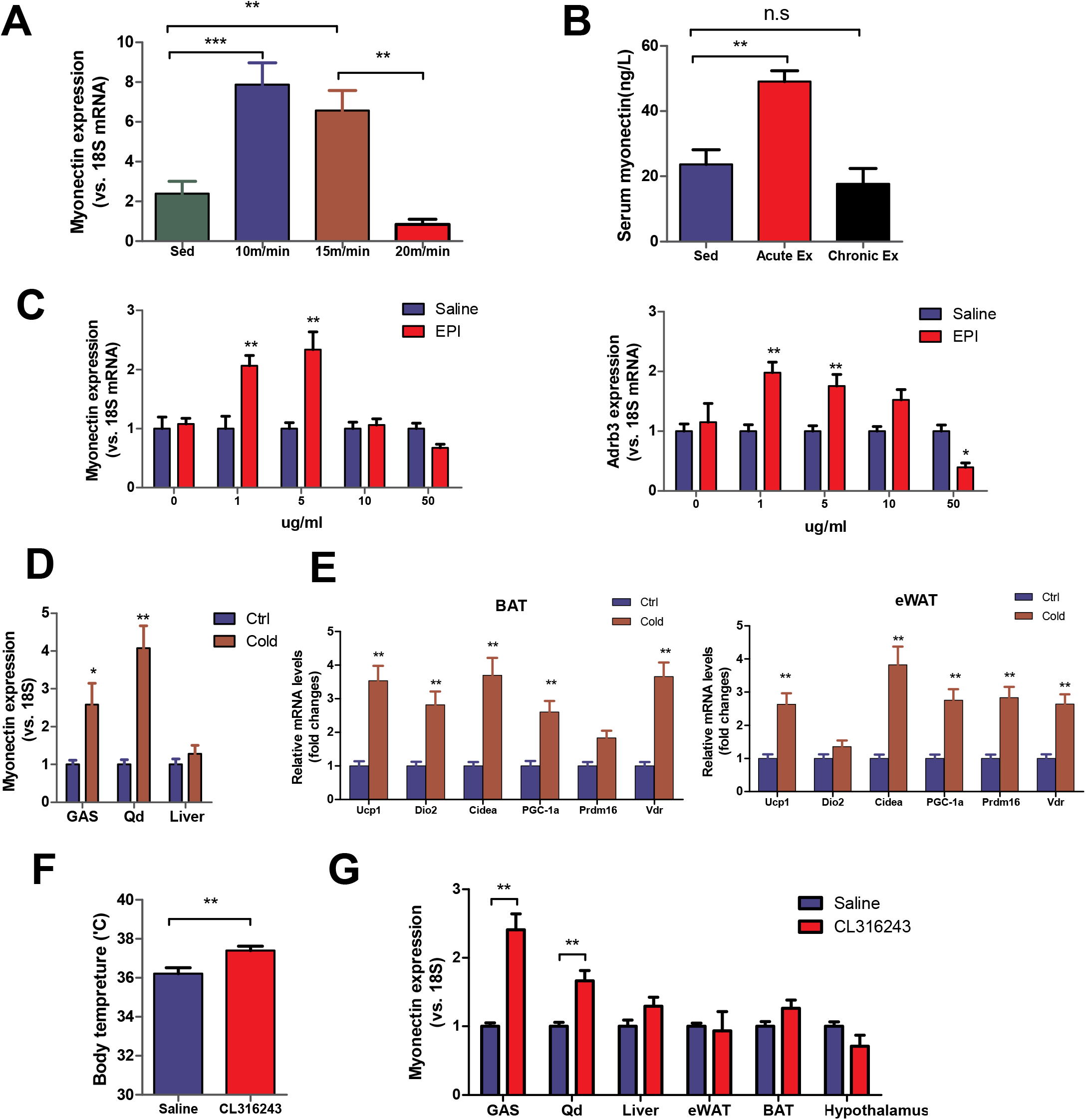
Physiological and pharmacological activation of sympathetic nerve system increases the expression of skeletal muscle Myn. (A) Myonectin expression in gastrocnemius muscle in C57BL/6J mice treated with one bout of treadmill running for 40min at the speed of 10, 15, or 20 m/min. Muscles were quickly removed and frozen for PCR immediately after exercise. n = 8/group. (B) Serum myonectin levels in mice with acute exercise (one bout of running at 13m/min for 40min) and chronic exercise (running for 3 weeks at 13m/min, 40min/day). n = 6–8/group. (C) Myonectin and Adrb3 expression in the cultured gastrocnemius muscles. n = 6/group at each concentration point of EPI. (D) Myonectin expression in gastrocnemius (GAS), quadriceps muscles (Qd) and liver after cold exposure. n = 8/group. (E) Gene expression of thermogenic markers in BAT and eWAT after cold exposure. Ucp1, uncoupling protein 1; Dio2, Deiodinase Type 2; Cidea, cell death-inducing DNA fragmentation factor-alpha-like effector A; PGC-1a, Peroxisome proliferator-activated receptor gamma coactivator-1 alpha; Prdm16, PR domain containing 16; Vdr, vitamin D receptor. n = 8/group. (F) Forehead temperature of mice treated with CL316243 in the morning. n = 8/group. (G) Myonectin expression in various tissues of mice after CL316243 treatment. n = 8/group. Statistical significance was evaluated by one-way ANOVA or unpaired two-tailed Student’s t test. Data are means ± SEM. Besides the indicated significance, *p < 0.05, **p < 0.01 versus Saline in Fig.3C, *p < 0.05, **p < 0.01 versus Ctrl in Fig.3D and 3E.

We next explore whether cold stress that often enhances sympathetic activity can increase Myn expression in skeletal muscle. Similarly, cold stress not only increases the expression of Myn in skeletal muscle (Fig. 3D), but also leads to the expression of thermogenic genes in BAT and WAT (Fig. 3E). In addition, we injected CL316243, a beta3-adrenergic receptor agonist that induces muscle hypertrophy and increased strength(Puzzo et al, 2016), into mice to mimic sympathetic activity. The treatment induced higher body temperature and enhanced Myn expression in skeletal muscle (Fig. 3F, G), suggesting that sympathetic activity increases Myn expression. As described above, our results indicate that both stress training and exercise increase Myn expression in skeletal muscle through sympathetic activity and adrenergic signaling.

### Myonectin attenuates glycemic response to EPI and activates insulin signaling in white and brown adipose tissue

To further determine the role of Myn in the diet-induced obesity, we designed adeno-associated viruses overexpressing Myn (AAV+) for skeletal muscle and treated mice by tail vein injection once a week during dietary intervention. At the 4^th^ week, Myn gene therapy significantly improved glucose intolerance and insulin resistance. At the same time, gene therapy reversed the hyperglycemic response to EPI induced by HFD (Fig. 4A). Within 30 minutes after EPI injection, blood glucose levels in the HFD group surged to nearly 20mmol/L, but Myn (AAV+) significantly attenuated this response (Fig. 4A, B). Compared with HFD-fed mice, Myn gene therapy not only increased serum Myn levels, but also reduced serum levels of leptin and triglyceride (Fig. 4C). However, there were no significant changes in serum glucose, insulin and glycerol levels (Fig. S3A). In addition, Myn gene therapy reduced epididymal and inguinal fat depot but not weight gain in HFD mice (Fig. S3B). Body composition analysis showed that Myn gene therapy reduced body fat mass (Fig. S3C). As we expected, Myn gene therapy increased Myn expression in the skeletal muscle, but had no effect in liver, adipose tissue and brain (Fig. S3D). These results suggest that Myn gene therapy targets skeletal muscle to protect against diet-induced obesity in mice.

**Fig.4.**
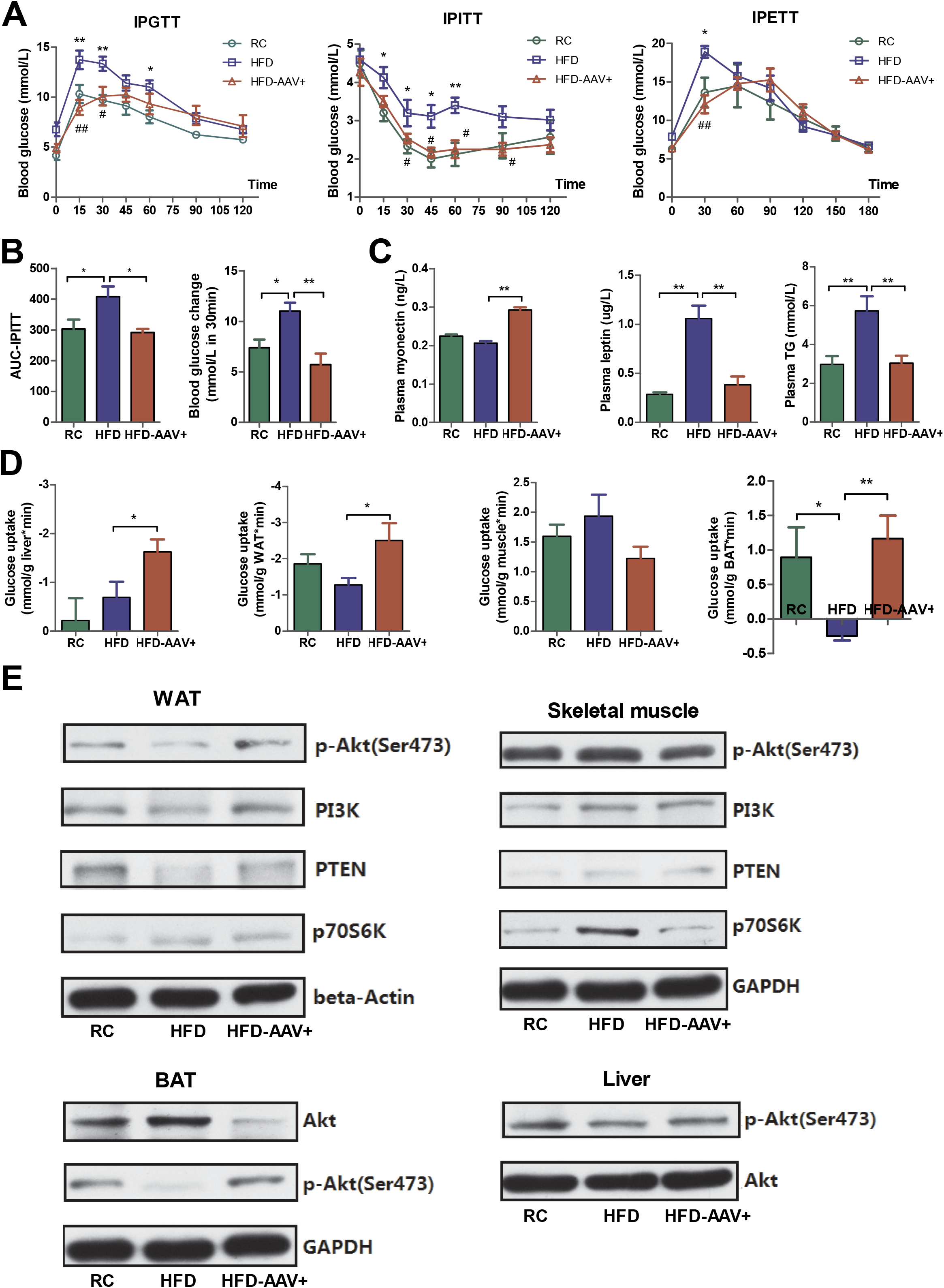
Myonectin gene therapy in HFD-fed mice reduces EPI sensitivity but activates insulin signaling in adipose tissues. (A) Glucose tolerance tests (IPGTT), Insulin tolerance tests (IPITT) and Epinephrine tolerance tests (IPETT) in Myn(AAV+)-treated mice. n = 7/group. (B) Area under curve (AUC) in the IPITT and blood glucose changes in 30 min after EPI injection in the IPETT. n = 6–7/group. (C) Plasma myonectin, leptin, and triglyceride (TG) levels. n = 10–12/group. (D) Insulin-stimulated glucose uptake in the cultured tissues of liver, eWAT, gastrocnemius and BAT. n = 6/group. (E) Western blot analysis of insulin signaling proteins in eWAT, BAT, skeletal muscle and liver. n = 4/group. Statistical significance was evaluated by one-way or two-way ANOVA with Tukey post hoc tests. Data are means ± SEM. Besides the indicated significance, *p < 0.05, **p < 0.01 versus RC, #p < 0.05, ##p < 0.01 versus HFD in Fig.4A.

We next explored the effect of Myn gene therapy on the insulin sensitivity in liver, skeletal muscle and adipose tissue. In the tissue culture incubated with insulin, Myn overexpression enhanced insulin-stimulated glucose uptake in liver and WAT, but not in skeletal muscle (Fig. 4D). It is noteworthy that HFD inhibited glucose uptake of BAT, but Myn gene therapy reversed this effect (Fig. 4D). Myn overexpression also increased Akt phosphorylation at Ser473 in both WAT and BAT (Fig. 4E, S3E), suggesting the activation of insulin signaling. The activation of insulin signaling in BAT was almost reversed, consistent with the insulin-stimulated glucose uptake (Fig. 4E). However, these results were not observed in skeletal muscle and liver (Fig. 4E, right). These data suggest that Myn gene therapy targeting skeletal muscle enhances the insulin signaling of WAT and BAT, thereby improving the metabolic status of mice with HFD.

### Myonectin inhibits HPA axis and yet promotes EPI-stimulated WAT lipolysis

In contrast to the increase in Myn levels, we found that Myn (AAV+) caused a decrease in serum levels of norepinephrine (NE), EPI(Fig. 5A) and corticosterone (Fig. 5B). Thus, we next explicitly tested the hypothesis that Myn suppresses HPA axis, resulting in reduced HFD-evoked HPA responses. HFD-fed mice have elevated serum corticosterone and ACTH compared with chow-fed controls (Fig. 5B). Notably, this HPA responses is consistent with previous studies in which mice or rats were exposed to high-fat diet or high-fat ketogenic diet(McNeilly et al, 2015; Ryan et al, 2018), suggesting that high-fat feeding enhances HPA axis tone in a manner that resembles sympathetic overdrive in chronic stress exposure. As expected, mice receiving AAV+ exhibited a rapid decrease in serum corticosterone and ACTH relative to HFD controls (Fig. 5B). Concurrently, mice receiving AAV+ exhibited inhibition of the HPA axis. In this case, plasma corticosterone were increased by the exposure to acute stress and exercise, and yet this increase was significantly attenuated among mice receiving AAV+ (Fig. 5C). Moreover, parameters of HPA functionality at the level of mRNA transcripts are reduced in hypothalamus of mice receiving AAV+ (Fig. 5D), implying an inhibition of HPA axis.

**Fig.5.**
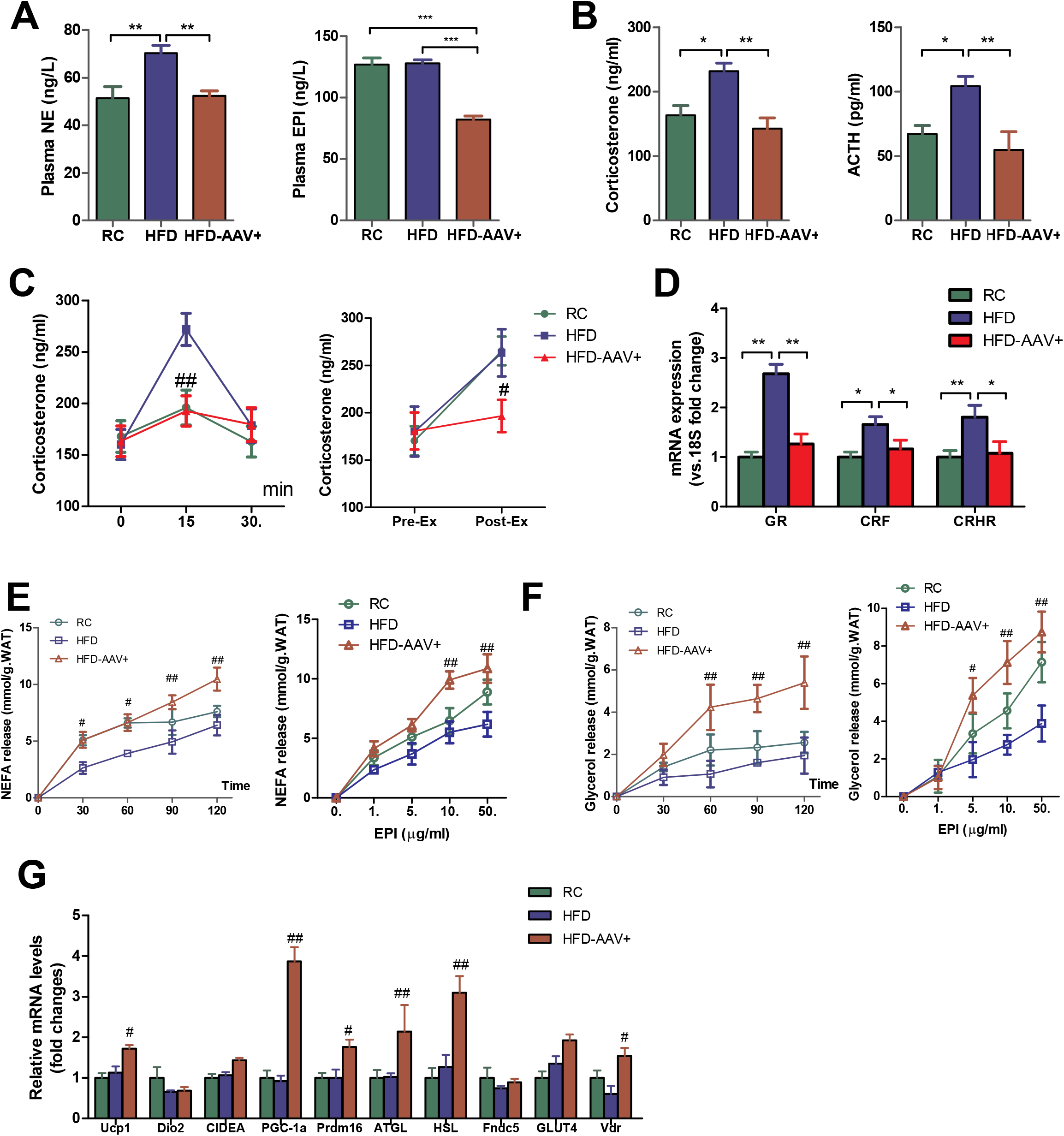
Myonectin inhibits HPA responses to HFD and yet promotes EPI-stimulated WAT lipolysis. (A) Basal level of plasma norepinephrine (NE) and epinephrine (EPI). n = 10 – 12/group. (B) Basal level of plasma corticosterone and ACTH. n = 10 – 12/group. (C) Changes in plasma corticosterone levels in mice exposed to acute restraint stress (left) and treadmill running (right). n = 6 – 7/group. (D) Gene expression of GR, CRF, and CRHR in hypothalamus. GR, Glucocorticoid Receptor; CRF, Corticotropin Releasing Factor; CRHR, Corticotropin Releasing Hormone Receptor. n = 8/group. (E) EPI-stimulated NEFA release from eWAT in the time- and dose-dependent test. n = 6/group at each time point (left), n = 6/group at each concentration point of EPI(right). (F) EPI-stimulated glycerol release from eWAT in the time- and dose-dependent test. n = 6/group at each time point (left), n = 6/group at each concentration point of EPI(right). (G) Gene expression of beige/brown adipocyte markers in the eWAT. Ucp1, uncoupling protein 1; Dio2, Deiodinase Type 2; Cidea, cell death-inducing DNA fragmentation factor-alpha-like effector A; PGC-1a, Peroxisome proliferator-activated receptor gamma coactivator-1 alpha; Prdm16, PR domain containing 16; ATGL, adipose triglyceride lipase; HSL, hormone-sensitive lipase; Fndc5, fibronectin type III domain containing 5; Glut4, glucose transporter 4; Vdr, vitamin D receptor. n = 8/group. Statistical significance was evaluated by one-way or two-way ANOVA with Tukey post hoc tests. Data are means ± SEM. Besides the indicated significance, *p < 0.05, **p < 0.01 versus RC, #p < 0.05, ##p < 0.01 versus HFD.

The activation of the HPA axis increased WAT lipolysis rates and hepatic acetyl-CoA content, which are essential to maintain gluconeogenesis during starvation(Perry et al, 2018). Therefore, we further explored whether Myn-induced HPA axis down-regulation affected lipid mobilization. Although Myn (AAV+) had no effect on serum glycerol levels (Fig. S3A), we found that Myn increased the sensitivity of eWAT to EPI. In the cultured adipose tissue, EPI resulted in a significant increase in NEFA release (p < 0.001 for variation with time; Fig. 5E, left). Moreover, EPI resulted in a dose-dependent increase in NEFA release (p < 0.001 for variation with EPI concentration; Fig. 5E, right). There were differences in the dynamics of the lipolytic responses to EPI between groups (p < 0.01 for site difference; Fig. 5E). Specifically, Myn (AAV+) promoted eWAT lipolysis stimulated by EPI. These site-specific differences in the lipolytic response were also shown in glycerol release (Fig. 5F). Furthermore, beige/brown adipocyte markers PGC-1a, Ucp1, Vdr, Prdm16 and hormone-sensitive lipase HSL and ATGL were upregulated in the eWAT after Myn (AAV+) treatment (Fig. 5G). Our data indicate that the lipolysis of WAT is enhanced by increased Myn levels independent of the HPA axis. Instead, Myn may inhibit the HPA axis in a negative feedback manner.

### Myonectin kncokdown enhances glycemic response to EPI and inhibits BAT thermogenesis

To further confirm the relationship between Myn and the EPI sensitivity of blood glucose, we designed adeno-associated viruses with Myn knockdown (AAV−) and transfected mice by tail vein injection. Contrary to our expectations, the knockdown of Myn also improves glucose tolerance. In HFD-fed mice, the blood glucose increased sharply 15 min after glucose injection and then decreased, but the Myn knockdown slowed down the increase in blood glucose (Fig. 6A). As expected, Myn knockdown increased the sensitivity of blood glucose to EPI. Mice were more sensitive to EPI after Myn (AAV−) treatment (Fig. 6A, B). Myn knockdown also down-regulated skeletal muscle Myn expression and reduced serum Myn levels (Fig. 6H). In addition, Myn knockdown increased serum insulin levels but decreased serum EPI, NE, and triglyceride levels (Fig. 6C, D). These results suggest that reducing Myn expression enhances glycemic response to EPI, which may predispose individuals to the risk of hyperglycemia during chronic stress or sympathetic activity.

**Fig.6.**
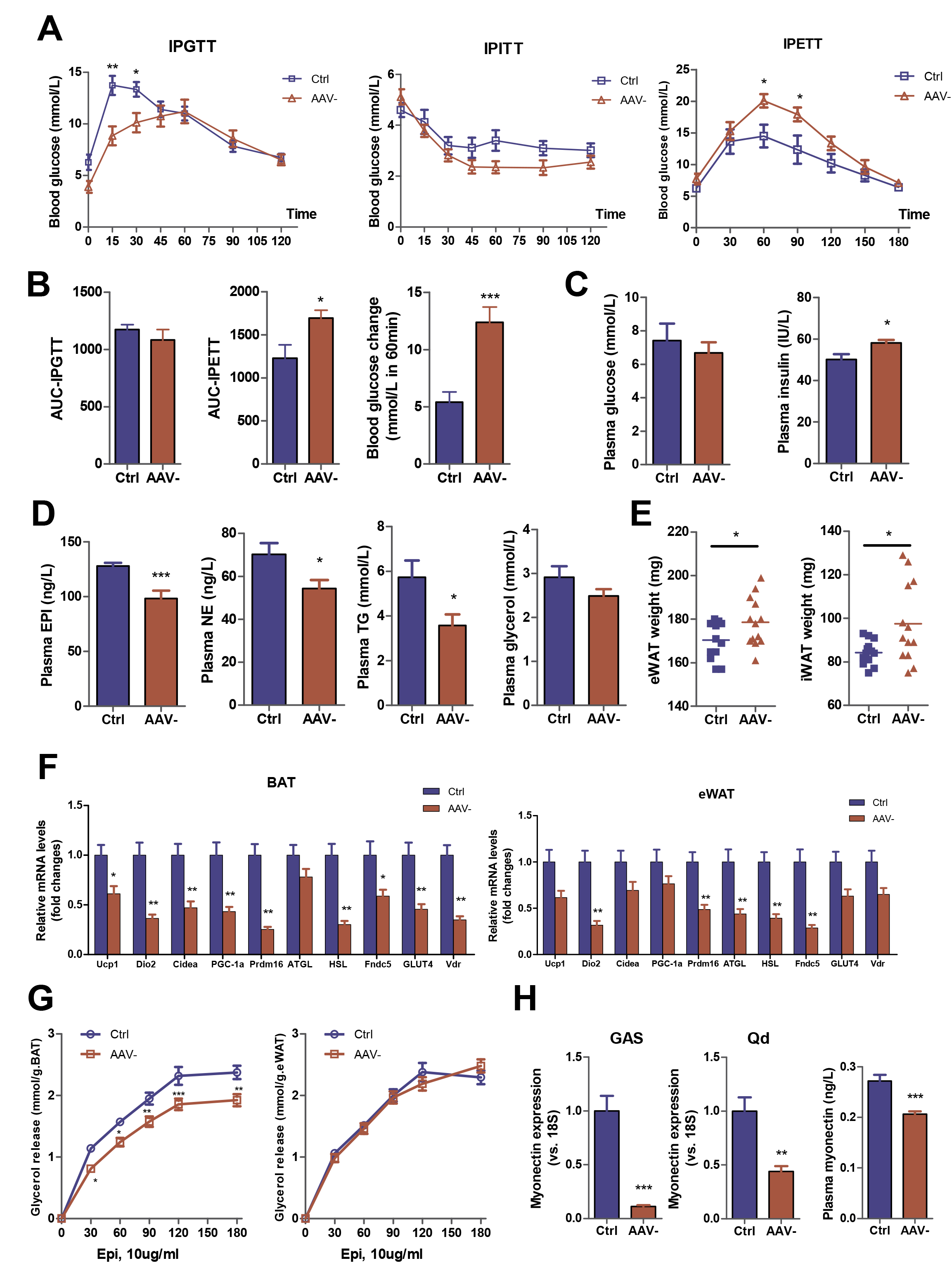
Reducing myonectin expression enhances the EPI sensitivity of blood glucose and yet inhibits BAT thermogenesis in HFD-fed mice. (A) Glucose tolerance tests (IPGTT), Insulin tolerance tests (IPITT) and Epinephrine tolerance tests (IPETT) in myonectin-AAV(−)-treated mice and control groups. n = 7/group. (B) Area under curve (AUC) in IPGTT and IPETT, and blood glucose changes in 60 min after EPI injection in the IPETT. n = 6–7/group. (C) Plasma glucose and insulin levels. n = 10–12/group. (D) Plasma epinephrine (EPI), norepinephrine (NE), triglyceride (TG) and glycerol levels. n = 10–12/group. (E) Epididymal and inguinal fat weight in mice with myonectin knockdown. n = 13/group. (F) Gene expression of thermogenic markers in BAT and WAT. Ucp1, uncoupling protein 1; Dio2, Deiodinase Type 2; Cidea, cell death-inducing DNA fragmentation factor-alpha-like effector A; PGC-1a, Peroxisome proliferator-activated receptor gamma coactivator-1 alpha; Prdm16, PR domain containing 16; ATGL, adipose triglyceride lipase; HSL, hormone-sensitive lipase; Fndc5, fibronectin type III domain containing 5; Glut4, glucose transporter 4; Vdr, vitamin D receptor. n = 8/group. (G) EPI-stimulated glycerol release from BAT and eWAT in 180 min after EPI was added into the cultured tissues. n = 6/group. (H) Myonectin expression in skeletal muscle and plasma myonectin levels in mice with myonectin knockdown. GAS, gastrocnemius muscle; Qd,quadriceps muscles. n = 8/group. Statistical significance was evaluated by unpaired two-tailed Student’s t test or two-way ANOVA with Tukey post hoc tests. Data are means ± SEM. *p < 0.05, **p < 0.01, ***p < 0.001 versus Ctrl.

In addition to the difference in glycemic response to EPI, we found that reducing Myn expression increased epididymal fat and inguinal fat weight (Fig. 6E). Mice receiving AAV− showed reduced thermogenic gene expression, such as Ucp1, Pgc1α, Dio2, Prdm16 and Cidea, in BAT (Fig. 6F). Indeed, Myn (AAV−) reduced EPI-induced lipolysis of BAT (Fig. 6G), consistent with reduced BAT thermogenesis. The EPI-induced lipolysis in WAT was not altered in mice receiving AAV− (Fig. 6G), although the mRNA levels of some thermogenic genes was decreased significantly (Fig. 6F). These data suggest that Myn controls glycemic response to EPI and regulates lipolysis and thermogenesis in brown adipose tissue.

### Myonectin kncokdown reverses the beneficial effects of stress training and exercise on glucose homeostasis

To address whether Myn is necessary for stress and exercise to improve glucose metabolism in HFD-fed mice, we treated mice with Myn AAV− during stress training and exercise intervention. Fig. 7A illustrates the study design. In brief, C57BL/6J mice were treated with HFD and AAV− injection for 4 weeks. Mice were treated with regular intermittent chronic stress (ICS) or treadmill running (Ex) for 3 weeks and then subjected to glucose, insulin and EPI tolerance test at the end of the 4^th^ week. Three weeks of stress training and running exercise significantly improved HFD-induced glucose intolerance, but Myn knockdown reversed this effect (Fig.7B, C, IPGTT). Three weeks of stress training and running exercise did not significantly improve insulin sensitivity, and reducing expression of Myn did not affect the insulin sensitivity of HFD mice (Fig. 7B, C, IPITT).

**Fig.7.**
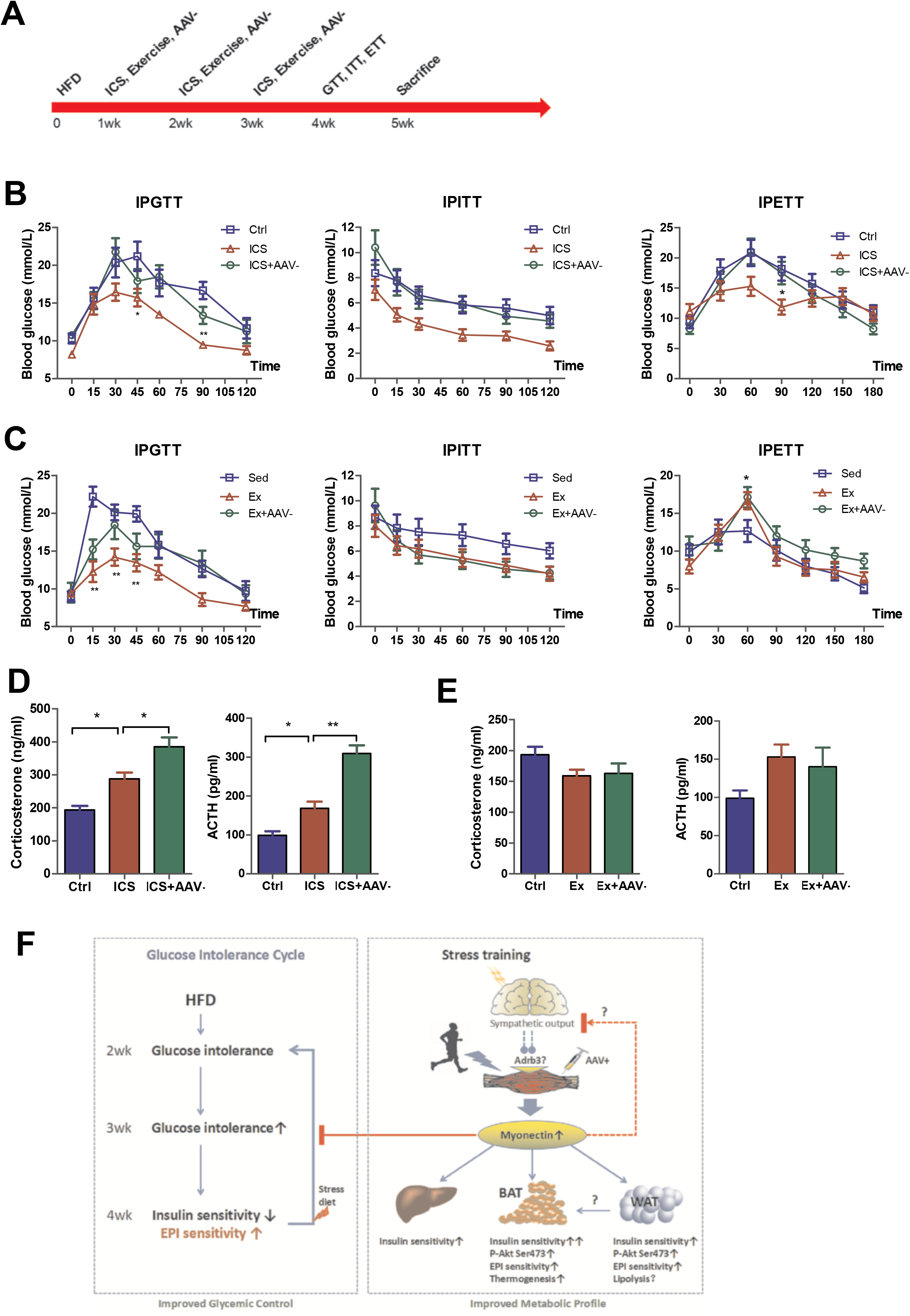
Reducing myonectin expression removes the beneficial effects of stress training and exercise on glucose homeostasis in HFD-fed mice. (A) Study design for the combination of myonectin knockdown with stress training (ICS, intermittent chronic stress) or regular exercise in HFD-fed mice. (B) Glucose tolerance tests (IPGTT), Insulin tolerance tests (IPITT) and Epinephrine tolerance tests (IPETT) in mice after stress training. n = 7/group. (C) Glucose tolerance tests (IPGTT), Insulin tolerance tests (IPITT) and Epinephrine tolerance tests (IPETT) in mice after running exercise training. n = 7-8/group. (D) Basal level of plasma corticosterone and ACTH in mice after stress training. n = 7/group. (E) Basal level of plasma corticosterone and ACTH in mice after running exercise training. n = 7-8/group. (F) Model depicting the role of myonectin in converting sympathetic output into the regulation of glucose intolerance and metabolic profiling. Statistical significance was evaluated by one-way or two-way ANOVA with Tukey post hoc tests. Data are means ± SEM. Besides the indicated significance, *p < 0.05, **p < 0.01 indicates ICS versus Ctrl in Fig.7B, and Ex versus Sed in Fig.7C, at the indicated time point.

IPETT showed that stress training reduced the sensitivity of blood glucose to EPI, but reducing Myn expression enhanced glycemic response to EPI (Fig. 7B, IPETT), suggesting the role of Myn in the glycemic response to EPI. However, the IPETT experiment showed that exercise further increased the glycemic response to EPI, and Myn knockdown could not reverse exercise effects (Fig. 7C, IPETT), indicating that physical exercise and stress training had differences in regulating EPI sensitivity of blood glucose. Studies have shown that chronic stress increased sympathetic nervous activity and elevated basal corticosterone levels and rapid glucocorticoid production (Lowrance et al, 2016). However, chronic exercise enhanced HPA-axis negative feedback and thus reduced HPA responses to stress or next bout of exercise (Hare et al, 2014). Thus, serum corticosterone and ACTH levels in response to stress or exercise training reflect the difference between stress and exercise (Fig. 7D, E). Together, our data indicate that Myn is essential for stress or exercise to improve glucose tolerance in HFD-fed mice.

## Discussion

Prior to this study, we have noticed that the important feature of childhood and adolescent obesity is sympathetic overactivity and in vivo resistance of lipolysis to EPI (Bougneres et al, 1997; Qi & Ding, 2016). Since sympathetic activation is intended to increase beta-adrenergic thermogenesis and expend excess energy, adipose tissue insensitivity to adrenergic stimulation in the young age leads to fat storage and even adiposity. However, sympathetic activation could be initiated in turn as a consequence of accumulating adiposity in order to prevent further fat storage in the development of obesity with aging (Seals & Bell, 2004). This suggests that increased sympathetic activity and fat storage constitute a vicious cycle that promotes each other in the development of obesity. The main reason for obesity is that adipose tissue is insensitive to adrenergic stimulation. Therefore, it is of great significance to explore the mediator to enhance the EPI sensitivity of adipose tissue for improving metabolic disorders and obesity.

The first important finding of this study is that increased glycemic response to EPI is a secondary event after glucose intolerance in the diet-induced obesity. Insulin resistance occurs after glucose intolerance when EPI sensitivity increases. Consistent with previous studies, HFD impairs the EPI sensitivity of adipose tissue (Gaidhu et al, 2010). Insulin resistance is characterized by an inability to reduce blood glucose and to promote glucose uptake in the tissues. Our results showed that EPI resistance was only found in the adipose tissue, and the EPI sensitivity of blood glucose increased instead. This suggests that HFD leads to increased sympathetic activity, consistent with previous study showing that HFD alters leptin sensitivity and elevates sympathetic nerve activity and arterial pressure in rabbits (Prior et al, 2010). Additionally, HFD increased heart rate and evoked metabolic disruptions (Simas et al, 2018), suggesting the elevated sympathetic activity in the diet-induced obesity. Increased EPI sensitivity subsequently may further aggravate glucose intolerance under stress or after diet, leading to the breakdown of glycemic control. Thus, glucose intolerance combined with increased EPI sensitivity in the early stage of HFD constitutes a vicious cycle that leads to worse glucose metabolism disorders eventually (Fig.1I).

Another major advance is that we identified Myn as a stress-responsive myokine that may break the vicious cycle and thus improve glucose tolerance. In order to avoid stress-induced depression in mice and try to reproduce the weight loss effect of stress, we modified the CUMS program by reducing stressed weeks and daily stress time. Stress training improved glucose tolerance and increased EPI sensitivity of white and brown adipose tissues in HFD mice. Notably, stress training resulted in an increase in skeletal muscle Myn expression and an increase in serum Myn levels. Previous studies have identified Myn as a muscle-derived myokine, which is upregulated by exercise and downregulated by HFD (Otaka et al, 2018; Seldin et al, 2012). We believe that Myn is also a potential mediator produced by the skeletal muscle as a psychological stress-responsive tissue. Our results demonstrated that, in the absence of stress training, exogenous AAV+ with Myn overexpression can alone effectively improve HFD-induced glucose intolerance, and increase the insulin and EPI sensitivity of BAT and WAT. In the presence of stress training, silencing Myn removes the beneficial effects of stress training on glucose intolerance. This suggests that Myn produced by skeletal muscle disrupts the vicious cycle between EPI sensitivity and glucose intolerance, thus glycemic control gets better (Fig.7F). However, it is essential for survival to maintain the sensitivity of the sympathetic adrenal system and thus to respond quickly to stress. This study showed that stress training improved glucose homeostasis and prevented diet-induced obesity at the expense of impaired EPI sensitivity. Further studies are needed to assess the risk of poor glycemic response to EPI due to psychological stress.

For adipose tissue, increased insulin sensitivity contributes to glucose uptake and utilization, and also contributes to lipogenesis and obesity, so it is of great significance for adipose tissue lipolysis and thermogenesis elicited by sympathetic innervation. Myn promotes fatty acid uptake in adipocytes and hepatocytes and thus improves systemic lipid homeostasis (Seldin et al, 2012). However, this study showed that stress training and Myn AAV+ enhanced lipolytic response to EPI in WAT and BAT and increased BAT thermogenesis. These results suggest that Myn plays an opposite role in regulating the EPI sensitivity of blood glucose and adipose tissue. It is the tissue-specific difference that improves glucose homeostasis and lipid metabolism reciprocally. However, excessive exercise intensity was also observed to inhibit Myn expression in skeletal muscle. In one hand, this reflects that skeletal muscles have resistance and self-protection against excessive exercise stress; on the other hand, increasing Myn expression may lead to stronger glycemic EPI resistance and higher EPI sensitivity of adipose tissue. As a result, the regulation of blood glucose and lipid metabolism will go to the negative side. Compared with exogenous stress, exercise is an endogenous way to enhance sympathetic activity. Although stress training improved glucose homeostasis, it impaired the EPI sensitivity of blood glucose. Exercise not only improved glucose homeostasis, but also protected the EPI sensitivity of blood glucose. Compared with stress training, exercise-induced endogenous sympathetic activity achieves the best coordination between glucose homeostasis and lipid metabolism through Myn.

It is equally important for human to maintain the stability of blood glucose and the sensitivity of blood glucose to EPI. Although studies have shown that hyperglycemia is harmful to multiple organs such as pancreas, kidney, brain, vessels and heart (Brownlee, 2013; Knudsen et al, 2019; Nagareddy et al, 2013), the elevated blood glucose under EPI stimulation is a necessary response to the activity of sympathetic-HPA axis (Peters et al, 2002; Rojas et al, 2015). This study proposed that the glucose intolerance cycle results from the combined effect of impaired insulin sensitivity and increased EPI sensitivity in the early stage of dietary obesity. Chronic stress can regulate glucose tolerance through the HPA axis, but the final blood glucose level is the result of the interaction between tissues and organs such as liver, muscle, fat, kidney and intestine (Aschbacher et al, 2014; Fu et al, 2009; Giudetti et al, 2019; Kovacevic et al, 2018; Sanghez et al, 2016). Skeletal muscle is the organ for human to escape invasion and capture food in human evolution. Changes in gene expression and atypical endocrine of skeletal muscle during strenuous exercise may provide neural feedback and regulate metabolism of other peripheral organs through myokines, and eventually make the best match between sympathetic nervous activity and the metabolic level of peripheral tissues (Baskin et al, 2015; Covington et al, 2016; Zhao & Karpac, 2017). Irisin as a serum factor produced by skeletal muscle mediates muscle-kidney crosstalk and suppresses metabolic reprograming and fibrogenesis during kidney disease (Peng et al, 2017). Our study proposes the following hypotheses: Myonectin is an important signal of skeletal muscle atypical endocrine, and its expression and release into serum are caused by various sympathetic activities (such as stress, exercise, cold exposure, beta-3 receptor agonists, etc). However, in the case of excessive exercise intensity, skeletal muscle may resist sympathetic overdrive, which is manifested by the inhibition of Myn expression in skeletal muscle during high-intensity exercise (Fig.3A). Myn released into blood enhances hepatic insulin sensitivity and adipocytes insulin sensitivity, lipolysis, and thermogenesis through some unknown receptors. Myn improves glycemic control through these metabolic reprogramming among peripheral tissues (Fig.7F). Exercise and stress training contribute to glucose homeostasis through Myn expression, however, only exercise maintains the flexibility of blood glucose.

These studies demonstrate that Myn functions as a stress-responsive target to improve glucose intolerance of HFD-fed mice and repress hyperglycemic response to EPI. Reciprocally, Myn enhances EPI-stimulated lipolysis of adipose tissue in vitro, which mediates the beneficial effects of stress training. Nevertheless, these studies have some limitations. First, the mechanism by which Myn inhibits glycemic response to EPI is inconclusive. Since IPETT was conducted after fasting, the reason for the increase of blood glucose should be glycogenolysis and gluconeogenesis stimulated by EPI. Myn promotes fatty acid uptake in hepatocytes (Seldin et al, 2012), partly support our result that Myn reduced plasma TG level in HFD mice (Fig. 4C) and thus potentially promotes lipid conversion in liver. Although Myn increased insulin-stimulated glucose uptake in the liver (Fig. 4D), further evidences in hepatic adrenergic signaling and gluconeogenesis are needed to explain why Myn inhibits the glycemic response to EPI. Second, we cannot rule out the independent effect of cold exposure and fasting. In this study, we emphasized that stress training is a program of neuropsychological stress to induce Myn expression. In fact, cold stress and fasting are among the stressors of this program, and they are randomly assigned to mice. Studies have shown that cold stress or intermittent fasting improves metabolism and glycemic control, promotes brown fat thermogenesis (Wang et al, 2013), and prevents obesity and so on (Crunkhorn, 2018; Liu et al, 2019; Peterson, 2019). Although our study supports stress training as an optional program to improve glucose metabolism and prevent from obesity, further studies are needed to explore the best way to increase sympathetic activity such as exercise, which not only inhibits hyperglycemic response to diet, and also maintains the sensitivity of glycemic response to stress as well.

In conclusion, this study expands our understanding of the reciprocal regulation of glycemic control and lipid homeostasis in adipose tissue in the development of obesity. In the early process of diet-induced obesity, glucose intolerance appears first and then leads to insulin resistance and increased EPI sensitivity, constituting a vicious cycle that progressively aggravates glucose intolerance. Stress training induces an increase in Myn expression and its circulating levels, which breaks the vicious cycle and improves glucose homeostasis at the cost of impaired EPI sensitivity. Myn increases WAT lipolysis and BAT thermogenesis, contributing to stress-induced weight loss. These data suggest that myonectin functions as a stress-responsive muscle-derived myokine that converts sympathetic output into metabolic signals targeting adipose tissues and so on. Future work should focus on identifying some myonectin receptors that localize in the adipose tissues, and understanding how muscle-derived myonectin links sympathetic activity to human metabolic diseases, obesity, and diabetes.

## Materials and Methods

### Animals

C57BL/6 mice were purchased from Shanghai SLAC laboratory Animal Co., Ltd (SLAC, China). All animal procedures were conducted under protocols approved by the animal experimentation committee of East China Normal University, China. Mice were housed and bred in a temperature-controlled room (22°C) on a 12h light/dark cycle in cages. Body weight was recorded twice a week. Food intake and water drink were recorded daily over 1 week. Food and drinking water were administered ad libitum, and the mice were fed a high-fat diet (HFD; 45 kcal% fat) or a regular chow (RC; 10 kcal% fat, both from SLAC, Shanghai China). Body composition was measured using NMR (MesoMR23-060H-I, NIUMAG).

### Chronic Mild Stress (CMS) Training

Animals were subjected to shortened and intermittent CMS protocols as reported previously (Tye et al, 2013). Mice in CMS group experienced one stressor during the day and a different stressor during the night for only 3 weeks before behavioral testing. Well-validated and approved standard stressors were randomly chosen from the following list so as to be unpredictable for mice: cage tilt on a 45° angle for 1 to 16 h; food deprivation for 12 to 16 h; white noise (by radio) for 1 to 16 h; strobe light illumination for 1 to 16 h; crowded housing (7 to 8 mice in a 10cm×10cm×10cm plastic box with air holes) for 1 to 3 h; individual housing (separating cage-mates into single-housing cages) for 1 to 16 h; light cycle (continuous illumination) for 24 to 36 h; dark cycle (continuous darkness) for 24 to 36 h; water deprivation for 12 to 16 h; cold water swimming for 5 minutes, damp bedding (200 ml water poured into sawdust bedding) for 12 to 16 h.

### Acute Restraint Stress, Cold Exposure and Drug Treatment

#### Acute Restraint Stress

Mice were physically restrained in a well-ventilated, 50 cm^3^ conical polypropylene tube for 30 min. Blood was collected from a small incision in the tip of the tail (time 0) and then at 15 and 30 min. **Cold Exposure.** Mice were subjected to ice cold water swimming for 10-15 minutes per day for 16 weeks. **Drug Treatment.** Mice were treated with i.p. injection of saline vehicle or CL316243 (1mg/kg/day) for 3 weeks.

### Exercise Protocols

**Acute exercise** was performed on a treadmill for 40 min running. Mice in group were subjected to an exercise program at the speed of 10, 15 or 20 m/min, which is respectively equal to 70%, 80% or 90% VO2max as measured previously in C57BL/6J mice (Schefer & Talan, 1996). Mice were sacrificed immediately after running, and blood was collected and tissues were extracted rapidly and frozen in liquid nitrogen. **Chronic exercise** (Regular running exercise) was performed on a treadmill, 40 min/day at 13m/min with no inclination (75% VO2max), 5 day/week for 3 weeks. Exercise was performed at the same time every day (4:00-6:00 p.m.). Mice were sacrificed at rest, 24h after the last session of running exercise.

### Recombinant AAV Vectors and In Vivo Administration

Recombinant AAV vectors of serotype 9 encoding Myn cDNA sequence under the control of the ubiquitous CMV promoter or Myn RNAi sequence were produced in HEK293T cells and purified by a CsCl-based gradient method as described previously (Jimenez et al, 2013). A non-coding plasmid was used to produce empty vectors for control mice. The mice received AAV vectors once a week by the tail vein (4.69×10^10^/mouse). The study diets were started concurrently with the AAV injections.

### Behavioral Testing

Except sucrose preference, a video computerized tracking system (DigBehav, Jiliang Co. Ltd., Shanghai, China) was used to record behaviors of the animals. All testing equipment was thoroughly cleaned between each session. These behavioral tests were performed as described in our previous study (Liu et al, 2018).

#### Sucrose preference test (SPT)

Briefly, 72 h before the test mice were trained to adapt 1% sucrose solution (w/v): two bottles of 1% sucrose solution were placed in each cage, and 24 h later 1% sucrose in one bottle was replaced with tap water for 24 h. After adaptation, mice were deprived of water and food for 24 h, followed by the sucrose preference test, in which mice housed in individual cages had free access to two bottles containing 200 ml of sucrose solution (1% w/v) and 200 ml of water, respectively. At the end of 24 h, the sucrose preference was calculated as a percentage of the consumed 1% sucrose solution relative to the total number of liquid intake.

#### Forced swim test (FST)

The swimming sessions were conducted by placing the mice in cylinders (30 cm height × 10 cm diameters) containing 25°C water 20 cm deep so that the mice could not support themselves by touching the bottom with their feet. The FST was conducted for 5 min and immobility time was recorded. Floating in the water without struggling and only making movements necessary to keep its head above the water were regarded as immobility.

#### Tail-suspension test (TST)

The test was performed as described in our previous study (Liu et al, 2018). Mice were individually suspended by the tail to a vertical bar on the top of a box (30 × 30 × 30 cm), with adhesive medical tape affixed 2 cm from the tip of the tail. The immobility time was recorded for a 5-min test session. In TST, immobility was defined as the absence of any limb or body movements except those caused by respiration.

#### Open field test (OFT)

The OFT is routinely used to study anxiety-like behaviors in mouse. In addition, the OFT is also employed to evaluate the effects of antidepressant treatment. Each mouse was placed in the center of the open field (30×30×30 cm chamber, with 16 holes in its floor) for 5 min in a quiet room after weighed. Parameters assessed were the number of poking into holes and the traveled distance and time in the center. Poking number can be considered as an exploratory parameter, and central distance and time can be used to measure locomotor activity.

### Glucose, Insulin and EPI Tolerance Test

Mice were deprived of food for 16 h and then subjected to glucose, insulin and EPI tolerance test (IPGTT, IPITT and IPETT). Blood was collected from a small incision in the tip of the tail (time 0) and then at different time-points after an i.p. injection of glucose (1g/kg), insulin (0.75U/kg), and EPI (0.1mg/kg). Blood glucose levels were measured using a glucometer (Accu-Check^®^ Active, Roche). Blood lactate and plasma glycerol were determined with enzymatic colorimetric assay following the manufacturers’ protocols (Nanjing Jianchen Biotech).

### Histology

Harvested tissues were fixed with 4% paraformaldehyde and then embedded with paraffin. Tissue sections were sliced and stained with haematoxylin and eosin (H&E). Slides were digitally scanned using an Aperio ScanScope System to produce high-resolution images.

### Tissues Culture ex vivo

Gastrocnemius muscles, liver tissues, BAT or WAT were extracted from mice and preincubated for 2 h at 30°C in Krebs-Ringer bicarbonate HEPES buffer (KRBH) containing (mM): 129 NaCl, 5 NaHCO_3_, 4.8 KCl, 1.2 KH_2_PO_4_, 1.2 MgSO_4_, 2.5 CaCl_2_, 2.8 glucose, 10 HEPES, and 0.1% BSA at pH 7.4. For ex vivo glucose uptake assay, tissues were incubated for 2 h with insulin 60 mU/mL followed by measurement of 2-deoxy-[^3^H]-glucose uptake as described previously (Cantley et al, 2014). For dose- dependent analysis of glycolysis and lipolysis in tissues ex vivo, tissues were incubated for 15 min with EPI at 0, 1, 5, 10, or 50μg/mL followed by measurement of lactic acid, glycerol or NEFA contents in culture medium. For time-dependent analysis of lipolysis in tissues ex vivo, tissues were incubated for 120-180 min with EPI at 10μg/mL followed by measurement of glycerol or NEFA in culture medium per 30 min (Gaidhu et al, 2010). Lactic acid, NEFA and glycerol contents were measured following the manufacturers’ protocols (Nanjing Jianchen Biotech).

### Serum Biochemistry Measurement

Serum samples were gathered from blood collected by cardiac puncture. Serum concentrations of myonectin, insulin, leptin, NE, EPI, corticosterone, and ACTH were measured by ELISA. Serum glucose, triglycerides (TG), glycerol were determined with enzymatic colorimetric assay following the manufacturers’ protocols (Nanjing Jianchen Biotech).

### Tissue Lipid Measurement

Tissue TAG was extracted using the method described previously (Vatner et al, 2015). Pieces of tissues were placed in preweighed glass vials and weighed. Tissues were homogenized in ice cold 2:1 chloroform : methanol, and lipids were extracted with shaking at room temperature for 3-4 h. Sulfuric acid was added to ~100 mM, and samples were vortexed and then centrifuged to achieve phase separation. The organic phase was then transferred to another preweighed vial (the lipid vial). The extraction medium in the lipid vial was evaporated, and TAG was measured using a Triglyceride Reagent Kit following the manufacturers’ protocols (Nanjing Jianchen Biotech).

### Quantitative PCR

Total RNA was isolated from tissues using TRIzol (Invitrogen), according to the manufacturer’s instructions. RNA was reverse-transcribed (cDNA Synthesis Kit, TOYOBO, Osaka, Japan). cDNA was amplified by real-time PCR in a total reaction volume of 20μl using SYBR Green Realtime PCR Master Mix (QPK-201; TOYOBO, Osaka, Japan). Real-time PCR reactions were cycled in StepOne™ Real-Time PCR System (Applied Biosystems). Target gene expression was normalized to housekeeping gene 18S or GAPDH and expressed as 2^−ΔΔct^ relative to the control group, respectively.

### Western blot

For frozen tissue, lysates were prepared using a RIPA lysis buffer (50mM Tris (pH7.4), 150mM NaCl, 2mM EDTA, 1% Nonidet P-40, 50mM NaF, 0.1% SDS and 0.5% sodium deoxycholate with PhosStop Phosphatase-Inhibitor Cocktail tablet and protease-inhibitor cocktail tablet (Roche). The homogenate was centrifuged at 4°C for 10 min at 14,000g and the supernatant was used for western blot analysis. The protein content of the supernatant was quantified using bicinchoninic acid reagents and BSA standards. Equal amounts of protein were separated using a polyacrylamide SDS-PAGE gel. After SDS-PAGE, proteins were transferred to PVDF membrane. The membrane was blocked for 1 h at room temperature followed by incubation overnight at 4°C with primary antibodies including myonectin (AVISCERA BIOSCIENCE), PI3K, PTEN, p70S6K, phospho-Akt (Ser473), beta-Actin or GAPDH (Cell Signaling). After overnight incubation, blots were incubated with HRP-conjugated secondary antibodies at a dilution of 1:5,000 for 1 h at room temperature. Bands were visualized by ECL plus according to the manufacturer’s instructions (Thermo Scientific) and quantified using Image lab software.

### QUANTIFICATION AND STATISTICAL ANALYSIS

Data are presented as means±SEM. Significance was assessed by Student’s t-tests and/or ANOVA with Tukey post hoc tests for multiple comparisons where appropriate. Statistical analysis were performed with GraphPad Prism, and differences were considered significant when p<0.05. Significances are indicated in the figures according to the following: *p < 0.05, **p < 0.01, and ***p < 0.001. Statistical parameters can be found in the figure legends.

## ACKNOWLEDGMENTS

We thank Dr. Li Rui, the director engineer of Genomeditech (shanghai) Co.,LTD, for designing and producing recombinant AAV vectors of serotype 9 encoding Myn cDNA sequence or RNAi sequence. We thank Dr. Wu J.L. and Dr. Liu Y.M., technician of the Instrumental Analysis Center of Shanghai Jiaotong University, for their expertise and conversations related to body composition analyses and metabolic assays. Supported by Natural Science Foundation of China grants 31871208 and 31300977, Shanghai Natural Science Foundation (18ZR1412000) and the Key Laboratory Construction Project of Adolescent Health Assessment and Exercise Intervention of Ministry of Education, China (No. 40500-541235-14203/004).

## AUTHOR CONTRIBUTIONS

Z.Q. and W.L. conceived, designed, performed, and interpreted experiments and made figures. J.X., X.X., J.L., W.L., L.L., Z.C., and Z.H. performed animal care, chronic stress, treadmill exercise training, behavioral analysis, tolerance tests, and real-time PCR, interpreted a part of experiments and made some figures. X.Z., Q.Z., X.L., L.C., B.J. performed animal care, acute exercise, treadmill exercise training, tolerance tests, Western blot and real-time PCR, and made some figures. S.D. participated in interpreting results and supervising the experimental plan. Z.Q. and W.L. wrote the manuscript.

---

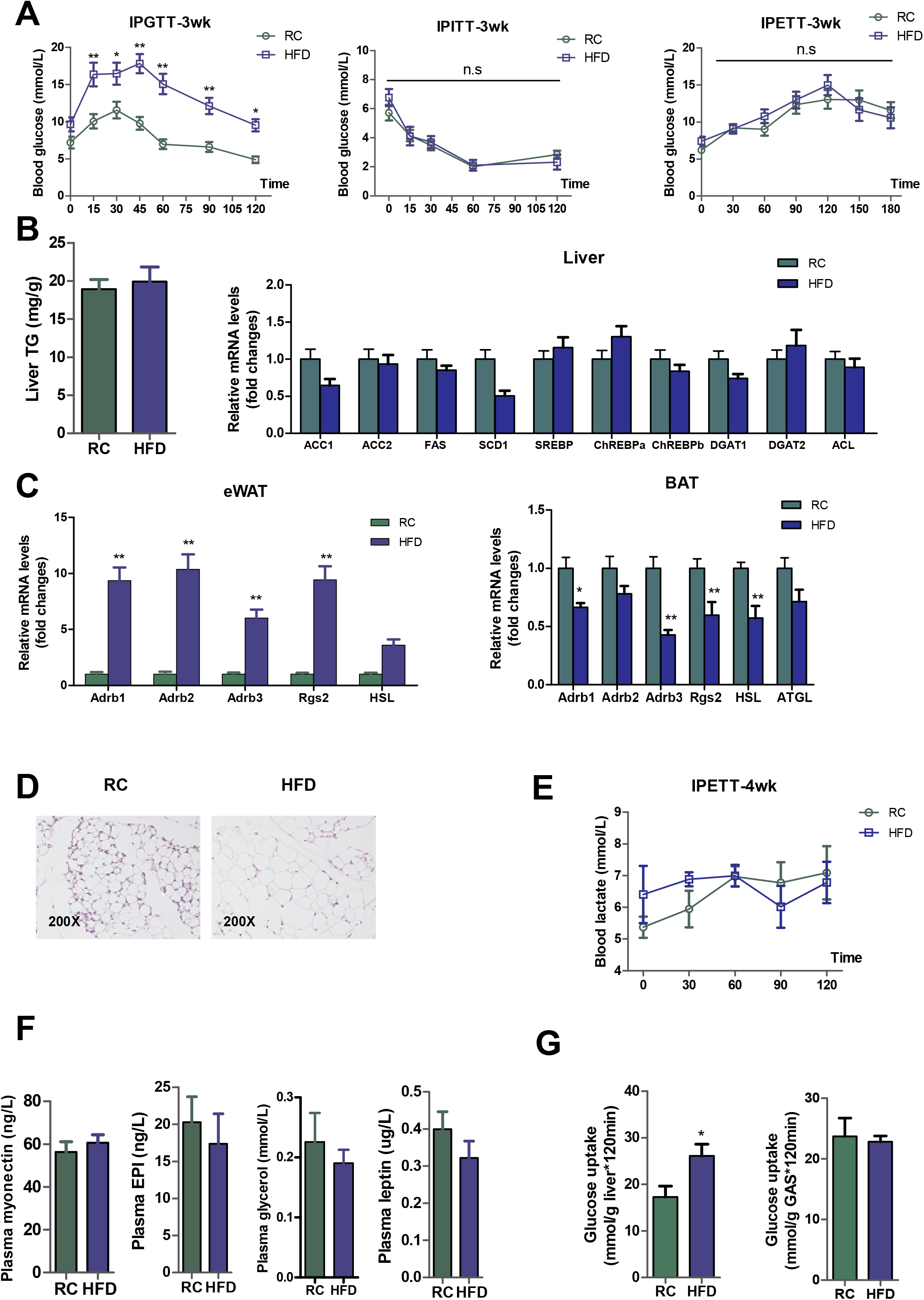

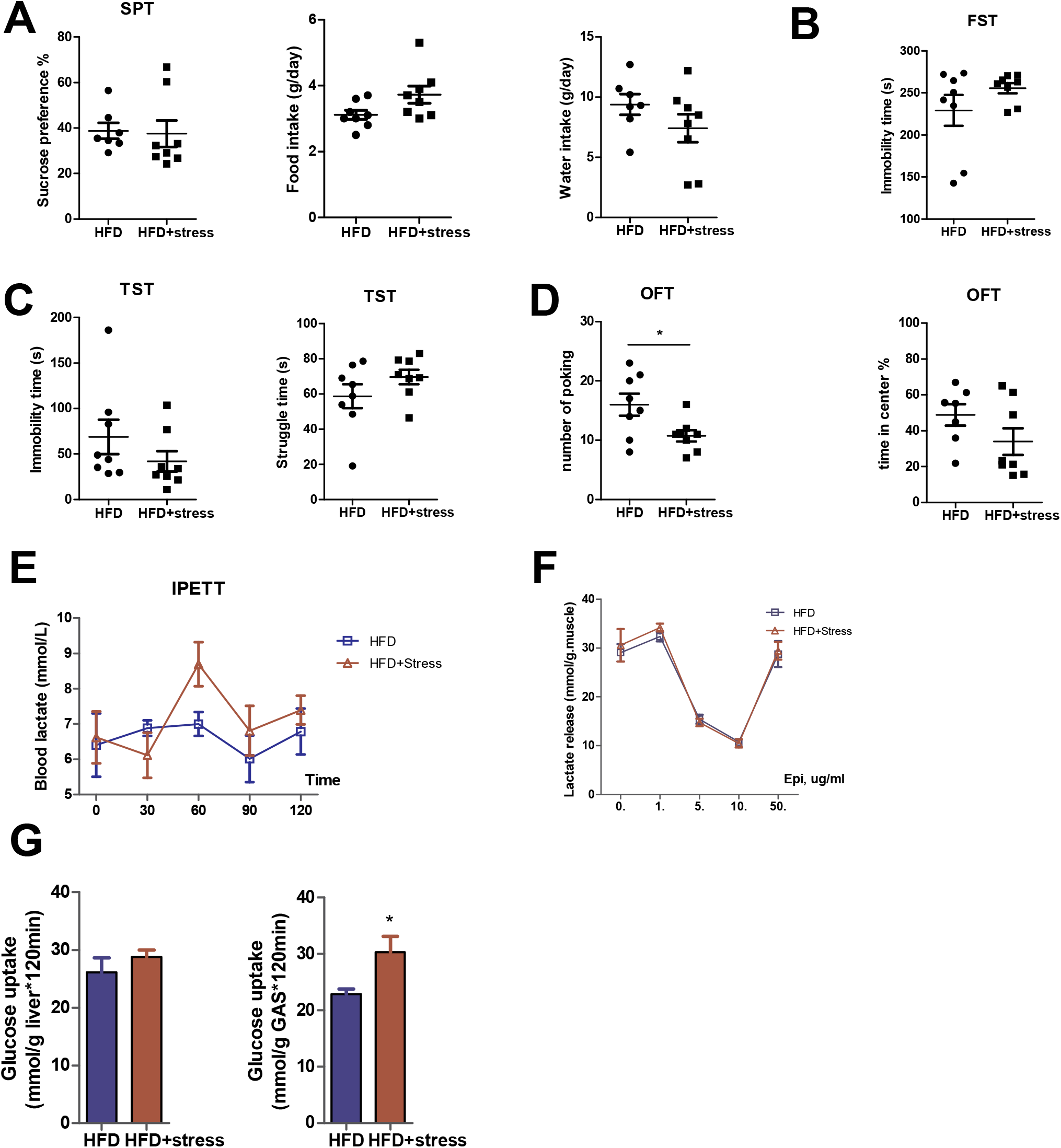

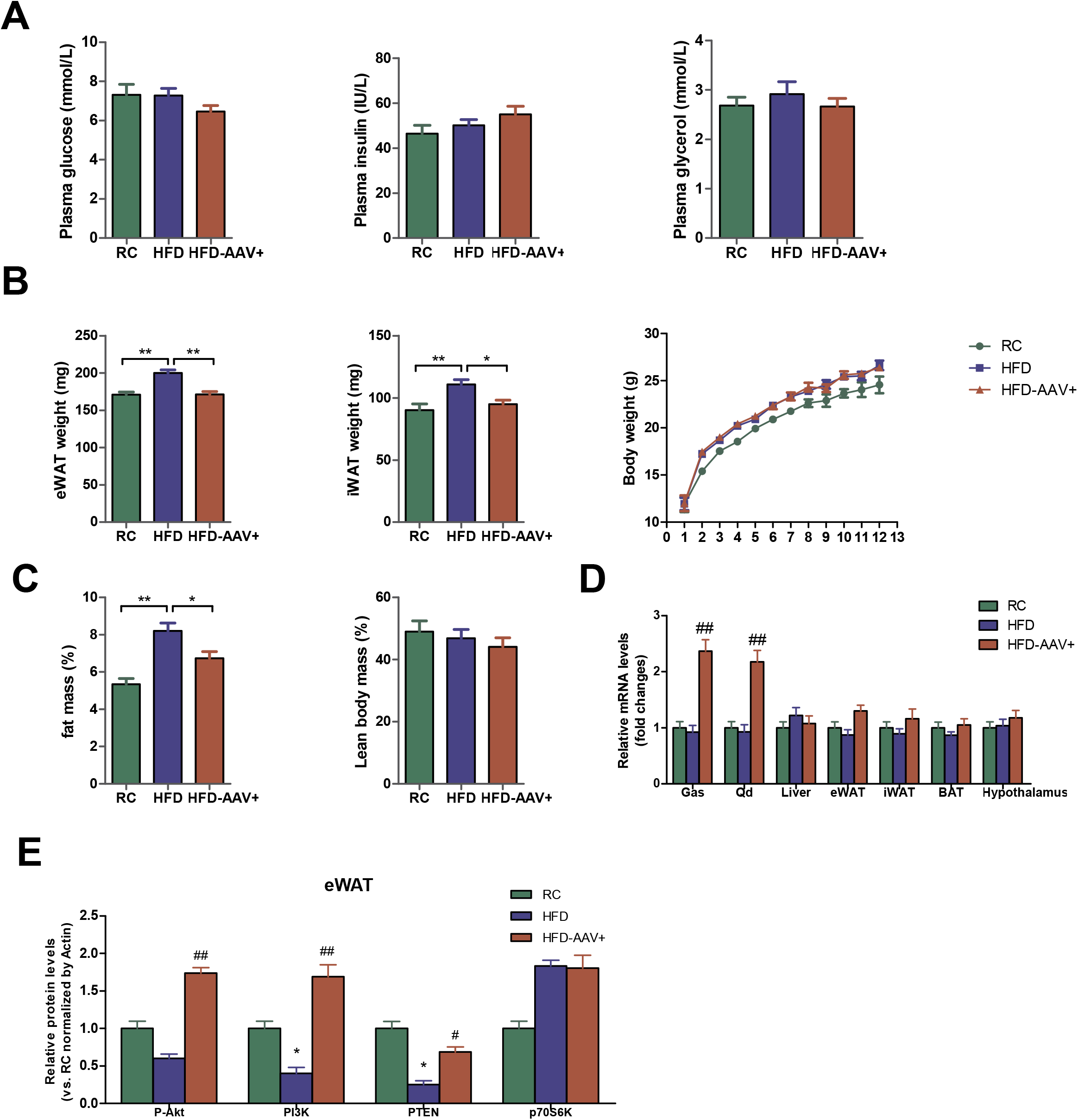

